# Selection on modifiers of genetic architecture under migration load

**DOI:** 10.1101/2021.12.24.474037

**Authors:** Stephen R. Proulx, Henrique Teotónio

## Abstract

Gene flow between populations adapting to differing local environmental conditions might be costly because individuals can disperse to habitats where their survival is low or because they can reproduce with locally maladapted individuals. The amount by which the mean relative population fitness is kept below one creates an opportunity for modifiers of the genetic architecture to spread due to selection. Prior work that separately considered modifiers changing dispersal, recombination rates, or altering dominance or epistasis, has typically focused on the direction of selection rather than its absolute magnitude. We here develop methods to determine the strength of selection on modifiers of the genetic architecture, including modifiers of the dispersal rate, in populations that have previously evolved local adaptation. We consider scenarios with up to five loci contributing to local adaptation and derive a new model for the deterministic spread of modifiers. We find that selection for modifiers of epistasis and dominance is stronger than selection for decreased recombination, and that selection for partial reductions in recombination are extremely weak, regardless of the number of loci contributing to local adaptation. The spread of modifiers that reduce dispersal depends on the number of loci, epistasis and extent of local adaptation in the ancestral population. We identify a novel effect, that modifiers of dominance are more strongly selected when they are unlinked to the locus that they modify. These findings help explain population differentiation and reproductive isolation and provide a benchmark to compare selection on modifiers under finite population sizes and demographic stochasticity.

**Author Summary:** When populations of a species are spread over different habitats the populations can adapt to their local conditions, provided dispersal between habitats is low enough. Natural selection allows the populations to maintain local adaptation, but dispersal and gene flow create a cost called the migration load. The migration load measures how much fitness is lost because of dispersal between different habitats, and also creates an opportunity for selection to act on the arrangement and interaction between genes that are involved in local adaptation. Modifier genes can spread in these linked populations and cause functional, local adaptation genes, to become more closely linked on a chromosome, or change the way that these genes are expressed so that the locally adapted gene copy becomes dominant. We modeled this process and found that selection on modifiers that create tighter linkage between locally adapted genes is generally weak, and modifiers that cause gene interactions are more strongly selected. Even after these gene interactions have begun to evolve, further selection for increased gene interaction is still strong. Our results show that populations are more likely to adapt to local conditions by evolving new gene interactions than by evolving tightly linked gene clusters.

## Introduction

Gene flow is a fundamental process that introduces, maintains, or reduces genetic variation between demes of a population adapted to local environmental conditions. Along with this variability comes a fitness cost, the “migration load”, that decreases the population mean fitness below that of a deme lacking immigration (Fisher, 1930; Haldane, 1937; Crow, 1958, 1970; Kondrashov, 1984). It has long been known that selection in natural populations can act to reduce the dispersal or recombination rates between loci experiencing recurrent mutation, dubbed the “reduction principle”, cf. (Altenberg et al., 2017). Much of the study on the reduction principle has been on proving that the direction of selection is towards reducing mixing of genotypes, rather than specifying the relative magnitude of selection on modifiers of the genetic architecture that can, in principle, diminish migration loads. Heritability for these modifiers has been described in the empirical literature including those that change dispersal rates (Saastamoinen et al., 2018), recombination rates (Rockman and Kruglyak, 2009; Penãlba and Wolf, 2020), and those that alter dominance (Durand et al., 2014; Billiard et al., 2021) and epistasis (Noble et al., 2017; Phillips, 2008) within and between locally adapted loci. Many of these types of modifiers may segregate concurrently within and between demes during and after the process of local adaptation. It is thus important to quantify the relative magnitude of potential selection on modifiers of genetic architecture to understand differentiation between populations, stability of hybrid zones and speciation.

Interest about selection on modifiers of genetic architecture under migration load has been spurred by the observation that genomic regions contributing to phenotypic differentiation and reproductive isolation between populations or incipient species do not often recombine (Turner et al., 2005; Nosil et al., 2009; Carneiro et al., 2013; Cruickshank and Hahn, 2014). How these “genomic islands” of differentiation and isolation originate and are maintained is intriguing. One possibility is that selection causes a build up of linked alleles that contribute to local adaptation, either by favoring mutations, such as chromosomal inversions, that eliminate recombination between pre-existing locally-adapted alleles (Kirkpatrick and Barton, 2006; Bürger and Akerman, 2011; Charlesworth and Barton, 2018), or by favoring mutations that just happen to be linked to locally-adapted alleles when they appear during ongoing local adaptation (Feder and Nosil, 2010; Feder et al., 2012; Flaxman et al., 2013; Yeaman, 2013; Aeschbacher and Bürger, 2014; Yeaman et al., 2016; Pontz and Bürger, 2022). Another possibility is that genomic islands of differentiation and isolation are created by genetic drift and the accumulation of mutations in genome locations of reduced recombination rates under allopatry, in a manner similar to how Dobzhansky-Muller incompatibilities arise and are maintained (Bank et al., 2012; Blanckaert and Hermisson, 2018).

Most theoretical work so far has described the expected direction and strength of selection on recombination modifiers under several simplifying assumptions (Kirkpatrick and Barton, 2006; Charlesworth and Barton, 2018; Bürger and Akerman, 2011; Akerman and Bürger, 2014). First, the recombination modifier is usually modeled as a chromosomal inversion so that its spread can be calculated without taking into account recombination of the modifier into alternative genotypes. Second, the fitness functions are generally assumed to be additive, meaning there is no dominance between locally-adapted alleles or epistasis between locally-adapted loci. Third, migration is usually assumed to be “continent to island” and thus dispersal is uni-directional. Under weak migration and no epistasis between loci, for example, Charlesworth and Barton (2018) show that the strength of selection on an inversion depends on the migration load, and that selection increases as the number of locally-adapted loci increase and is higher when the ancestral genotype has free recombination. When the modifier creates chromosomal inversions, the strength of selection on it is completely determined by the fitness load present in the structured population (Proulx and Phillips, 2005).

A separate literature has considered the evolution of dominance modifiers, dating back to early work by Fisher (1928) and Wright (1929). A major objection to Fisher’s notion that dominance modifiers could be selected to favor wild-type alleles under mutation-selection balance is that heterozygotes are rare and dominance modifiers are therefore only weekly selected (Feldman and Karlin, 1971; Karlin and McGregor, 1974; Wagner and Bürger, 1985). However, if other processes maintain heterozygotes at high frequencies, then selection for dominance modifiers can be substantial (Otto and Bourget, 1999; Van Dooren, 1999). For example, when there is dispersal between demes facing different environmental conditions, migration-selection balance can lead to the long-term maintenance of heterozygotes and result in selection for dominance modifiers (Otto and Bourget, 1999). Understanding selection on modifiers of epistasis is complex, depending on the sign of linkage disequilibrium between deleterious alleles being purged or beneficial alleles being favored with the mutant modifier alleles, as well as on the dominance interactions at each locus (Phillips et al., 2000; Pontz and Bürger, 2022). We do know, however, that in general an epistatic modifier affecting the total fitness load of a structured population is expected to be strongly selected relative to modifiers affecting only specific loci (Proulx and Phillips, 2005).

When there is multi-directional dispersal between demes, calculation of relative selection strength on modifiers of genetic architecture depends on the full distribution of genotype densities in the demes. Much of the discussion has centered on the distribution of offspring genotypes of a given parent, but this view obscures the fact that modifiers alter their own associations with alternative genotypes, and therefore that the strength of selection depends both on the frequency of genotype or environment backgrounds in the absence of the modifier as well as on the reproductive value of those states. For example, Proulx and Servedio (2009) explore this interaction when alleles affecting mating preferences are selected under migration-selection balance. They find that selection on changes in the transition probability from a particular genotype or environmental background will be weak if the frequency of that state in the stable stage distribution (absent the modifier) is low, even if the potential fitness benefit of a transmission modification is high. Conversely, selection on changes in the transition probability from a genotype or environment background into a different genotype or environment background will be weak if the absolute reproductive value of the new background state (absent the modifier) is low.

Selection on modifiers of the genetic architecture under migration load will be strongest for changes in the transition probabilities that affect relatively frequent states and cause a large change in the reproductive value of their descendant states, possibly leading to local extinction (Szep et al., 2021). Because modifiers affect these transition probabilities without altering the total number of offspring produced, any increase in the probability of transition between two states must be associated with an equal reduction of transmission to other states. Therefore, one must calculate the cumulative effect of these changes in transmission in order to quantify the magnitude of selection on modifiers of genetic architecture under migration load.

We here investigate selection in a scenario with bi-directional dispersal between two demes that face two habitats with differing environmental conditions, where a diploid genotype is determined by a linear chromosome with arbitrary recombination between several adjacent loci (up to five), and where fitness can include dominance and epistasis. Although we assume a linear chromosomal recombination map, if the recombination probability is 0.5, then we recover a scenario with freely segregating chromosomes. Mating between individuals, precisely syngamy, is assumed to be random. We numerically solve for the equilibrium – a steady-state between migration, selection, segregation, recombination and syngamy – of genotype frequencies in each deme and calculate the migration load. We then introduce a rare mutant modifier of the genetic architecture and calculate the selection strength on the invading modifier as the eigenvalue of the transition matrix. The modifier can independently alter the overall magnitude of the migration load, the dominance at single or multiple loci, the epistasis between loci, the recombination rate between loci, and the migration rate between demes.

## Methods

### Life-cycle

We consider two demes with reciprocal migration, each adapting to habitats with different environmental conditions. After birth, diploid individuals migrate between the two demes and are subject to viability selection within each deme (see next section for the description of the fitness function). After selection, gametes are produced by meiosis with Mendelian segregation and cross-over recombination between homologous chromosomes. Recombination is modeled as if there were a genetic linkage map of a single linear chromosome. The locations of the *k* loci are given by their index and a recombination vector describes the probability of crossing-over between successive loci, giving *k* − 1 recombination rates. After recombination, gametes are produced by defining a table of the frequency of all possible gamete types given each possible pair of parental haplotypes. Following (Kirkpatrick et al., 2002), a square matrix *G* with dimensions 2^*k*^ is defined such that each element *G*_*i,j*_ is itself a vector of length 2^*k*^ (technically *G* is a tensor of 3 dimensions). Each position *l* in the vector represents the probability that a parent with haplotypes *i* and *j* would produce a gamete with index *l*. The gamete production table is computed for situations with up to *k* = 5 local adaptation loci and *1* modifier loci. This model has a total of 6 loci with 64 haplotypes, and thus, the gamete production table has 64^3^ = 262, 144 entries. These gamete tables are defined generically without inserting numerical values and stored as an object. Values can be efficiently substituted into these stored objects, meaning that the gamete table does not need to be re-calculated during simulations, but it does need to be evaluated with the recombination parameter values once per simulation of the modifier invasion (see below).

The life cycle just described can be concisely expressed as:

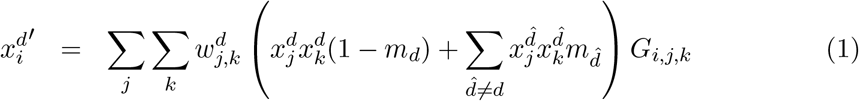

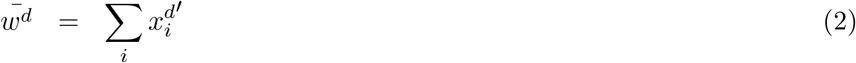

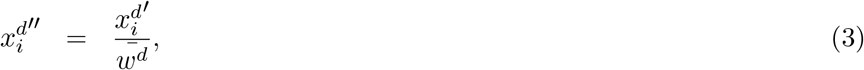

**Table 1:** Notation

| Variable | Definition |
| --- | --- |
| $k$ | The number of genes contributing to local adaptation |
| $l_i(h_1, h_2)$ | The effective number of locally adapted alleles |
| $S$ | Is the total fitness cost associated with the least-adapted genotype |
| $m_d$ | The migration rate from deme $d$ |
| $r_m$ | The recombination rate for the modifier |
| $r_i$ | The recombination frequency between locus $i$ and $i + 1$ |
| $\phi$ | The epistasis parameter in favor of the locally adapted allele in deme 1 |
| $\theta_i$ | The dominance parameter for locus $i$ in favor of the locally adapted allele in deme 1 |
| $G$ | The matrix storing the probability of producing each possible gamete haplotype for each diploid genotype. |
| $w_{ij}^d$ | The fitness of an adult carrying haplotypes $i$ and $j$ in deme $d$. |
| $\bar{w}^d$ | The mean fitness in deme $d$ |
| $x_i^d$ | The frequency of haplotype $i$ in deme $d$. |
| $\hat{z}$ | The parameters associated with the modifier allele are indicated with $\hat{\phantom{z}}$ |
| $\mathbf{A}$ | The invasion matrix for the mutant modifier allele. |
| $\lambda$ | The dominant eigenvalue of $\mathbf{A}$ |
| $L_d$ | The fitness load in deme $d$ |

where 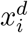 is the frequency of haplotype *i* in deme *d*, 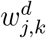 is the fitness of genotype (*j, k*) in deme *d, G*_*i,j,k*_ represents the probability that an adult with haplotypes *j* and *k* will produce a gamete of haplotype *i*, and 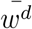 is the average fitness within deme *d*. Haplotype frequency in the next generation is given by 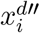.

We also have developed a notation system to describe the life-cycle and the dynamics of modifiers using linear algebra operators, and this is detailed in appendix A. The main benefit of the linear operator notation is that it separates the life cycle into modules that can be easily modified or amended, for instance to include non-random mating, parental effects, or non-Mendelian segregation.

### Fitness function

Diploid individual fitness is a function of both habitat and genotype. We model fitness by assuming that each locus has a parameter determining dominance between alleles at each of the *k* local adaptation loci, and that epistasis between all *k* local adaptation loci are determined by a single parameter. Our approach is to use generalized logistic functions to describe the *effective* number of locally adapted alleles. Specifically, for a population with *k* loci, each individual has 2*k* alleles that are each either locally adapted (allele state of 1 in deme 1) or not (allele state of 0 in deme 1). Fitness in habitat *d* for an individual that inherited haplotypes *h*_1_ and *h*_2_ is then:

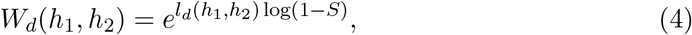

where *S* represents the total fitness cost of having the most maladapted genotype for the resident haplotypes, and *l*_*d*_(*h*_*1*_, *h*_2_) represents the effective fraction of alleles that are maladapted carried by an (*h*_1_, *h*_2_) individual in habitat *d*. In our parameterization, when the dominance and epistasis parameters are set to 0, *l*_*d*_(*h*_1_, *h*_2_) is equal to the actual fraction of alleles that are maladapted. In other words, when there is no epistasis or dominance the effective number of locally adapted alleles is the same as the actual number of locally-adapted alleles. If *l*_*d*_ = 1, then fitness is 1 − *S*, and if *l*_*d*_ = 0 then fitness is 1.

For dominance, each locus is parameterized by *θ*_*i*_, which determines how the alleles interact. At a locus *i*, the effective fraction of locally adapted alleles is 0 if the individual is homozygous for the maladapted allele, 1 if the individual is homozygous for the locally adapted allele. If the individual is heterozygous then locus specific fitness is *w*_*i*_ = 1 − *h*_*i*_*s* where

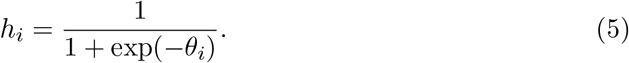

This gives *h*_*i*_ = 1/2 if *θ*_*i*_ = 0, indicating that if there is no dominance in either direction, and heterozygotes behave as if 1/2 of their alleles are locally adapted. In the limit as θ_i_ → ∞ the fraction adapted goes to 1, and as *θ*_*i*_ → −∞ the fraction adapted goes to 0.

Once we have calculated dominance at each individual locus, we sum across loci to determine an aggregate effect:

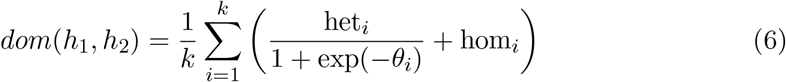

with het_*i*_ an indicator variable that takes on 1 if the individual is heterozygous at locus *i* and 0 otherwise, hom_*i*_ an indicator variable that takes on the value 0 if the individual is homozygous for the locally favored allele at locus i and 1 otherwise.

Epistasis is applied in a similar way as dominance except now the fraction of adapted alleles is calculated across loci:

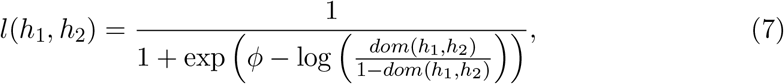

where *ϕ* is the epistasis parameter. When *ϕ* = 0, then *l*(*h*_1_, *h*_2_) = *dom*(*h*_1_, *h*_2_), so that the fraction of adapted alleles is unaltered by epistasis. The epistasis calculation depends on both *ϕ* and the fraction of adapted alleles determined by dominance. If dom(*h*_1_, *h*_2_) = 0, then *l* = 1 for any finite value of *ϕ*. Likewise, if dom(*h*_1_, *h*_2_) = 1, then *l* = 0 for any finite value of *ϕ*. If 0 < dom(*h*_1_, *h*_2_) < 1 then as *ϕ* → ∞ l approaches 0 while as *ϕ* → −∞ *l* approaches 1.

Under this formulation, fitness is additive when *ϕ* = 0 and *θ*_*i*_ = 0 (for all *i*). Further, changes in the *θ* and *ϕ* from 0 have symmetric effects on the log scale. As *ϕ* and *θ* move away from 0, the effect quickly saturates (Figure 1A). For dominance, if gene regulation is environment dependent, for example, one might find that a single modifier allele causes locally adapted alleles to become dominant in the habitats they perform best in and conversely more recessive in the habitats they perform less well (Figure 1B). This is reminiscent of classical physiological explanations of partial dominance (Kacser and Burns, 1981). Similarly for epistasis (Phillips et al., 2000), adding locally adapted homozygous loci has diminishing fitness returns in habitats where the population performs well, while it is synergistic in habitats where the population performs less well (Figure 1C).

**Figure 1:**
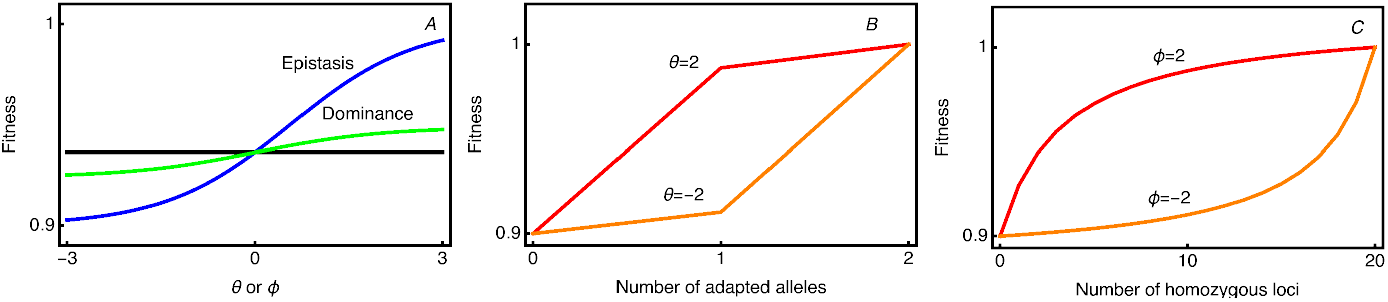
The fitness function showing the effects of dominance and epistasis. The total fitness cost is set at *S* = 0.1. Panel A shows the effect of changing the dominance and epistasis parameters in a genotype composed of 20 loci, 5 of which are homozygous for the adapted allele, and 10 of which are homozygous for the maladapted allele. The black line is the fitness value with no dominance or epistasis shown for reference. Note that in this example *θ* is the same for all loci. In panel B, the effect of dominance on fitness is shown for a single locus, as the number of adapted alleles is varied from 0 (homozygote) to 1 (heterozygote) to 2 (homozygote). The effect is shown for two values of the dominance parameter *θ*. Panel C shows the effect of the epistasis parameter in a genotype with 20 loci, as the number of loci homozygous for the adapted allele is varied. Two values of the epistasis parameter, *ϕ*, are shown.

### Ancestral population

We use iterations of the recursion equations shown above for the life-cycle to obtain a population of “resident” haplotypes at an equilibrium between standing variation, dispersal, recombination, mating and selection. We initialize the recursion equations with the locally favored haplotype being common (frequency of 0.9999) and with the other haplotype frequencies assigned at random. This avoids the pitfall of having the simulation maintain a symmetric but unstable equilibrium. We find equilibrium haplotype frequencies by iterating the recursion equations while censusing the population at the haploid gametic stage. Iteration occurs for a maximum of 5000 generations or until the haplotype frequencies change by less than 10^−8^ (see supplemental Mathematica S1 file for further details).

### Selection on the modifier mutant allele

The primary goal of our study is to calculate the magnitude of the leading eigenvalue of an invasion matrix describing modifiers of the genetic architecture of the resident population. The genetic architecture is fixed in the resident ancestral population, which means, for example, that the recombination rates between loci do not depend on the allele effects at those loci. Local adaptation can be maintained so long as dispersal is not too high and when selection for locally adapted alleles is relatively balanced among the habitats. Even when there is initially no dominance or epistasis determining fitness, heterozygosity is maintained at migration-selection balance as well as linkage disequilibrium between loci within and between demes. Implicitly, indirect selection on the modifier allele is created by linkage between the invading modifier allele and locally-adapted haplotypes and thus depends on the extent of heterozygosity in the resident population. In all of the analyses presented here we assume symmetric migration rates and selection coefficients, as this ensures that the fully polymorphic equilibrium is stable (Akerman and Bürger, 2014). Our method works under asymmetry as well, but it then requires additional checking to ensure that a polymorphism in local adaptation is maintained.

The invasion matrix for the mutant modifier allele, **A**, defines the number and genotype of mutant offspring produced per mutant individual in the prior generation. The construction of this matrix follows the same life-cycle as for the resident population, but differs in that the frequency of potential mates is set by the frequency of resident genotypes at the resident equilibrium, and of course in that the mutant parameters are applied. Thus, the matrix elements *A*_*i,j*_ represent the number of diploid adults in the following generation that carry the modifier and the mutant haplotype and resident haplotype denoted by the index *j* that descend from a single individual with the mutant haplotype and resident haplotype denoted by the index *i*.

The modifier is always placed in position 1 of the genetic linkage map. Each pair of adjacent loci experiences an independent chance of crossing-over given by the recombination vector. The modifier may alter the parameters which are indicated by Ŝ, 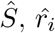 (recombination modifier), 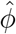 (epistasis modifier), 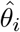 (dominance modifier), and 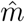 (dispersal modifier). Here, we consider modifier alleles that alter only one parameter at a time, although in principle the modifier may alter several or all the parameters at once. Because the mutant modifier allele is rare, matings between two individuals carrying it are even rarer. The recombination mutant modifier allele is dominant, all other mutant modifier alleles are co-dominant.

We start by indexing the vector describing the density of mutants as ν, which has dimension 2^2*k*+1^. This dimensionality comes from enumerating all combinations of mutant and resident haplotypes in both demes (2^*k*^ ∗ 2^*k*^ ∗ 2). The matrix **A** is then constructed in three stages, first with a matrix **D** combining the segregation and recombination matrix from the two demes and then applying migration, and third applying the matrix **F** to adults after migration. The matrix **R**^*j*^ has dimension 2^*k*^, and details how segregation, recombination and syngamy produce offspring in the next generation. If we define the component haplotypes of genotype *i* as *i*_*μ*_ and *i*_*r*_ for the mutant and resident haplotypes found in the diploid with index value *i*, then:

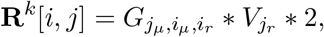

where 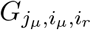 represents the probability that a gamete with haplotype *j*_*μ*_ is produced by a diploid bearing haplotypes *i*_*μ*_ and *i*_*r*_. This is multiplied by 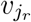 to represent the probability that the gamete fuses with a resident gamete bearing haplotype *j*_*r*_. The factor of 2 accounts for the fact that, at population dynamic equilibrium, each diploid adult produces an average of two successful gametes.

Migration then moves mutant genotypes between the two demes. For this, we create a block matrix composed of the recombination/syngamy matrices:

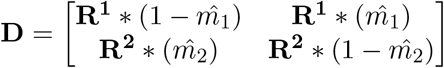

which has dimension 2^2k+1^.

After migration, we apply selection by creating a vector f of relative fitness values, for each diploid genotype. This vector has length 2^2k+1^ and its elements are:

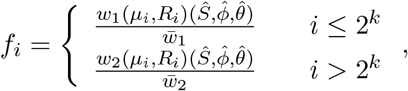

where the fitness function uses the mutant fitness parameters. Because the fitness function is applied at the adult stage, the transmission of genotypes in **D** to the next stage depends only on their column index, not their row index. We define a matrix *F*_*i,j*_ = *f*_*j*_

The transition matrix for the invasion modifier is then

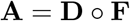

The invasion recursion for the modifier across generations *t* can be described by

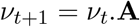

The selection coefficient of the mutant modifier allele, or long-term growth rate multiplier, is the dominant eigenvalue of **A**. With our approach, the potential loss by genetic drift of the mutant modifier is not modelled and thus we assume that once the modifier appears it will be established in the resident population and follow deterministic dynamics (Chelo et al., 2013). However, the invasion matrix generated here can be used to define a multi-type branching process to study stochastic establishment and will be considered in a future work.

## Results and Discussion

### Invading the ancestral population

We first examine selection on modifiers of the genetic architecture of an ancestral population characterized by multiplicative fitness and free recombination (i.e. *r*_*i*_ = 0.5, *ϕ*_*j*_ = 0, and *θ*_*i,j*_ = 0). We consider modifiers that alter one aspect of genetic architecture at a time (i.e. that change one of the fitness/recombination/migration parameters). We also assume that modifiers have symmetric effects in the two demes, such that *ϕ*_2_ = −*ϕ*_1_ and *θ*_*i*,2_ = −*θ*_*i,1*_. Biologically, this amounts to assuming that gene expression is environment independent and that fitness effects are opposite but symmetric between loci (Rice, 2002; Grieshop et al., 2021). Other assumptions are plausible, and may even be expected when gene expression is modified in an environmentally plastic fashion. Here we stick with this simple assumption of opposite effects and note that this is a conservative assumption with respect to the strength of selection on dominance modifiers. We set relatively high levels of prior local adaptation, with the total fitness cost of the most maladapted genotype being 0.1 and migration rates of 0.05. We further modeled complete linkage between the modifier locus and the first local adaptation locus (*r*_*m*_ = 0) and identical values for effect of a modifier on dominance at each locus 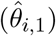 and recombination parameter 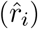. In all figures, modifier parameters are expressed relative to the maximum change possible for that parameter. For example, for recombination, 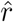 can be reduced from 0.5 down to 0, so we use the transformation 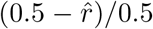 to get the fractional change. For dominance and epistasis, the effect of negative values is completely symmetric to that of positive values, representing the opposite polarity for the deme in which the locally adapted allele is dominant. The values of *ϕ* and *θ* are limited to a maximum of 3 because the effect of the modifier on the effective number of locally adapted alleles is saturated by this point (Figure 1). We used this maximum value of 3 to rescale the dominance and epistasis parameters for the figures.

Results of this analysis are shown in Figure 2. We plot the theoretical maximum for a modifier that does not directly alter fitness, which is strictly a function of the migration load (Proulx and Phillips, 2005). We also plot, for reference, the magnitude of selection on a modifier that reduces the total negative effect of maladapation, i.e. a modifier that reduces *S* directly. We find that modifiers of recombination rate have the smallest selection coefficients, and that there is very weak selection unless they almost completely eliminate recombination between locally adapted alleles. While selection for reduced recombination is slightly higher when more loci contribute to local adaptation, it is still the case that selection only becomes noticeably strong as recombination gets close to zero.

**Figure 2:**
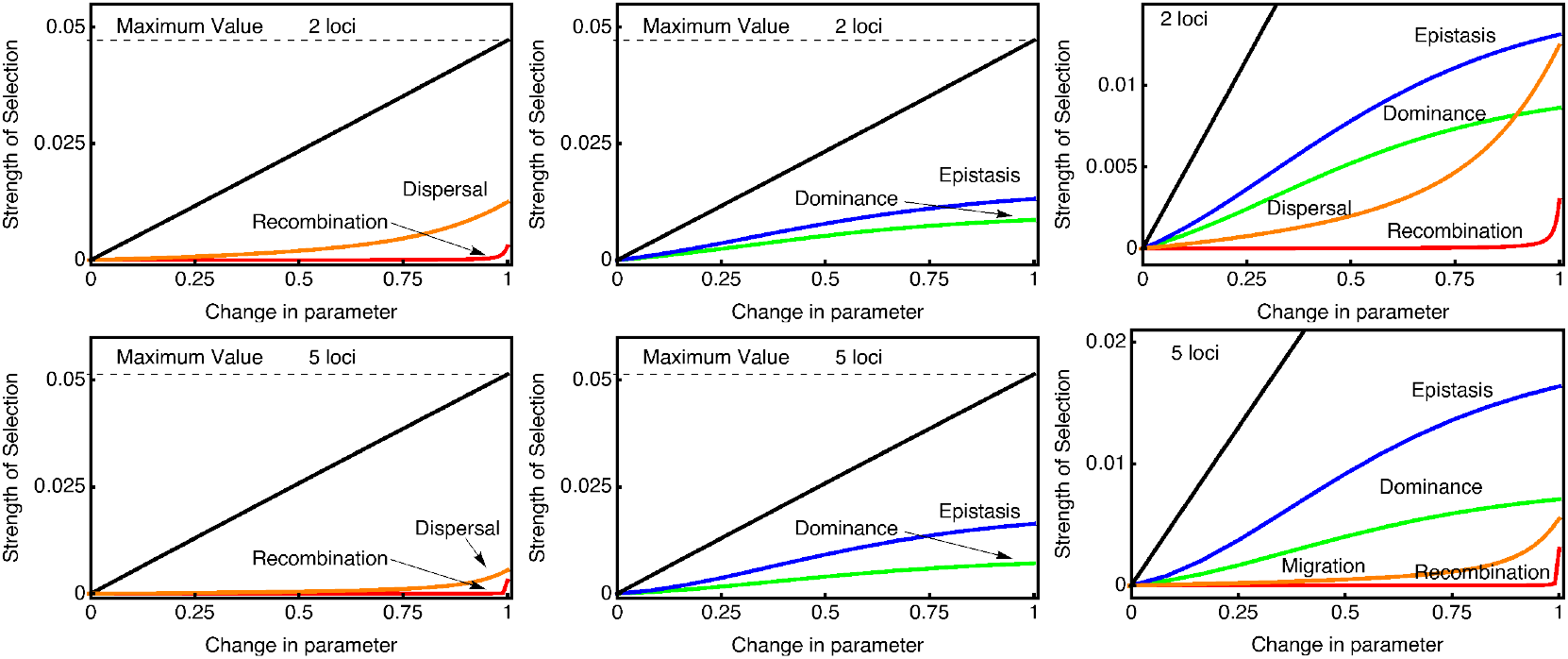
Initial strength of selection on different modifiers as measured by λ − 1. Parameter values for the ancestral population are *S* = 0.1, *M* = 0.05, *R* = 0.5; *θ* = 0; *ϕ* = 0. We plot the fraction of the maximum allowable change in the parameter on the x-axis. The dashed horizontal line represents the theoretical maximum for the spread of a modifier, while the solid black line shows the strength of selection for a modifier that reduces the total effect of the maladaptation (i.e. a modifier that reduces *S*). Top panels left to right show results for two local adaptation loci, while bottom panels left to right show results for five local adaptation loci. For dominance modifiers, the modifier is assumed to affect dominance at all local adaptation loci simultaneously. Likewise, the recombination modifier is assumed to reduce recombination between all loci simultaneously. We only show positive changes in dominance and epistasis, because negative values have identical effects but indicate that deme 2’s locally adapted allele is instead dominant.

Selection for reduced dispersal into unfavorable habitats is generally stronger than selection for reduced recombination. The strength of selection on migration modifiers decreases substantially as the number of loci increase, even though the magnitude of the migration load is slightly higher. This is because the migration modifier is directly linked to only a single local adaptation locus and can recombine into a more favorable genetic background after migrating into an unfavorable habitat.

Selection is strong on both epistasis and dominance, and saturates as the modifier parameter increases. Even small increases in the dominance or epistasis parameter results in an appreciable selection coefficient. Selection on epistasis is stronger than selection on dominance, for the same level of phenotypic bias, and this difference becomes more noticeable as more loci contribute to local adaptation. Overall, selection for modifiers of epistasis and dominance is stronger than selection for modifiers of recombination and dispersal.

### The number of local adaptation loci

We next varied the number of loci contributing to local adaptation while keeping the magnitude of dispersal and the migration load constant. More than 5 loci contributing to local adaptation becomes computationally straining, though asymptotic patterns can be observed as we go up to 5 loci. We determine the maximum possible strength of selection when one of the parameters is set to an extreme value. For recombination and dispersal modifiers this involves setting the parameter to 0. For epistasis and dominance modifiers it involves setting parameters to 10 because this causes complete dominance or epistasis.

Results show that as the number of local adaptation loci increases from 2 to 5 the total migration load goes up by about 10%, which increases the maximum potential strength of selection (Figure 3A). Selection on epistasis and recombination modifiers show increases of similar amounts, indicating that the fraction of the migration load that they can ameliorate remains nearly constant. Selection on dominance decreases only slightly with increase number of loci. Most different is the response of dispersal, where the strength of selection decreases by about 50%.

**Figure 3:**
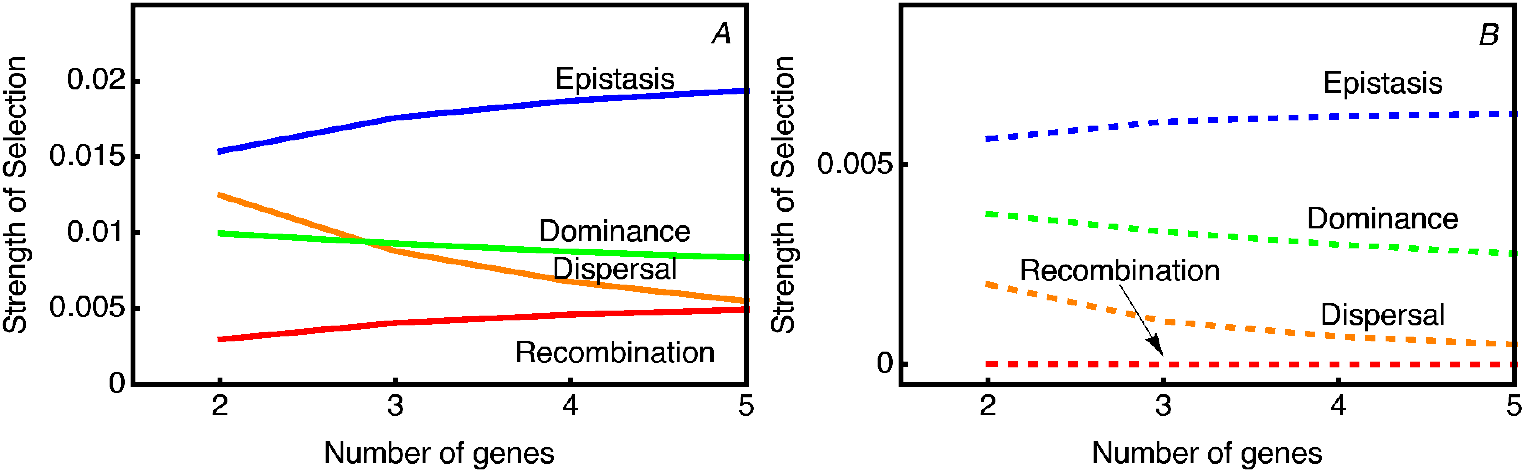
Strength of selection on different modifiers as measured by λ − 1. Parameter values for the ancestral population are *S* = 0.1, *M* = 0.05, *R* = 0.5; *θ* = 0; *ϕ* = 0. Panel (A) shows the strength of selection of the maximum effect for each parameter (for dominance and epistasis we use a parameter value of 10, for migration and recombination we use a value of 0). Panel (B) shows the effect when the parameters have a modest value (for dominance and epistasis we use a parameter value of 1.5, for migration and recombination we use a value of half the resident population value).

We also evaluated the effect of fitness loci on the strength of selection for moderate effect mutant modifier alleles. For dispersal and recombination modifiers we reduce their parameter values to 50% of the ancestral population rate. For epistasis and dominance we set the parameter to 1.1, because this leads to a 50% reduction in the fitness cost when half the alleles are locally adapted. We find the pattern of change is largely the same, with the main difference being that the strength of selection on the recombination modifier is relatively smaller than on the other modifier types (Figure 3B).

### Dominance modifiers

In Figure 2 results are presented for modifiers of dominance that affect all fitness loci simultaneously. It is thus interesting to consider modifiers of dominance that affect only one of the local adaptation loci. For this, the modifier is assumed to be at one end of a linear chromosome and completely linked (r=0) with the first local adaptation locus (called locus 1). In Figure 4 we show the analysis for a scenario where the ancestral population has free recombination (r=0.5) between successive fitness loci. When the modifier affects locus 1, the strength of selection is lower than when the modifier affects the second locus 2 or a higher index locus in scenarios with more than two loci.

**Figure 4:**
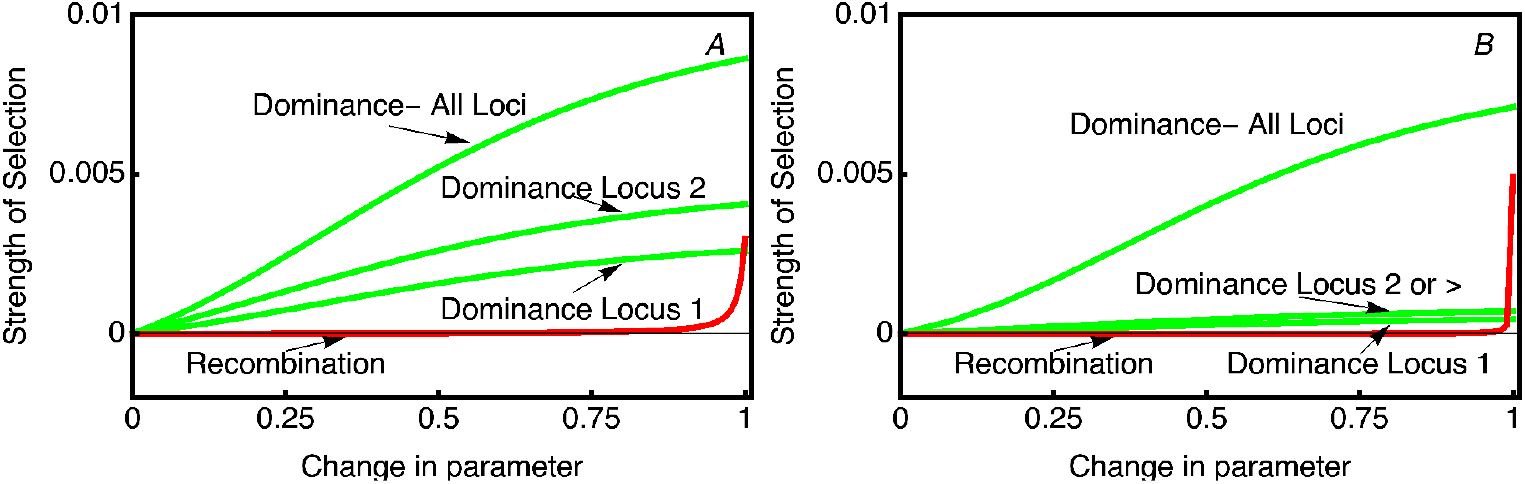
: Strength of selection on dominance modifiers is dependant on modifier linkage. Parameter values for the ancestral population are *S* = 0.1, *M* = 0.05, *R* = 0.5; *θ* = 0; *ϕ* = 0. In Panel (A) 2 loci contribute to local adaptation. The green curves show selection on a modifier that simultaneously affects dominance of all loci, or a modifier that affects dominance of the locus tightly linked to the modifier (locus 1) or a locus that freely recombines with the modifier (locus 2). In panel (B) 5 loci contribute to local adaptation. Loci 2-5 all freely recombine with the modifier, and so selection on modifiers that affect dominance at any of these loci have the same strength of selection.

As shown before in Figure 3, selection on modifiers affecting all fitness loci simultane-ously decreases slightly as the number of loci increases. When considering modifiers that affect dominance at single local adaptation loci, we find that the strength of selection is substantially lower, and that the effect of modifying multiple loci is more than additive. Regardless of the number of loci, the strength of selection is stronger when the modifier affects unlinked genes. Furthermore, when only two loci contribute to local adaptation the effect of position is much larger than when five loci contribute to local adaptation. Selection on dominance modifiers depends on two components, the frequency of heterozygotes in the ancestral population and the change in relative fitness that is caused by the modifier. If there is little potential effect of the modifier, i.e. because the cost of having the maladapted allele is low, then selection on the modifier will be weak. Likewise, if heterozygotes are at low frequency, i.e. because there is little migration and strong selection against mutants, then selection on the modifier will be weak. Neither factor alone is particularly predictive of modifier selection strength. For example, if we compare scenarios with different values of *S*, increases in *S* cause heterozygosity to decrease but cause selection on the modifier to increase (not shown).

### Ancestral levels of epistasis

While the heterozygosity of the ancestral population will determine selection on dominance modifiers, ancestral levels of linkage disequilibrium will determine selection on other modifiers of genetic architecture. In part, the ancestral population genetic equilibrium is a function of epistasis and therefore we can vary it to model how selection on the modifiers would change with an evolving genetic architecture, and not, as our modeling presupposes, that selection during the invasion is the same as when the modifier reaches intermediate frequencies.

Because we must consider a modifier of epistasis whose fitness effects are positive in one habitat and negative in the other it means that the maximum rate of spread of an epistatic modifier allele causing its bearer to have relative fitness of 1 is given by a migration rate dependent eigenvalue, λ(*A*), where:

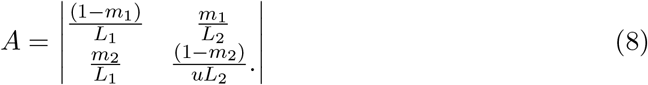

The theoretical maximum strength of selection for an epistasis modifier is then given by the eigenvalue of this matrix minus 1: λ(*A*) − 1 (Appendix B).

As the amount of epistasis increases, λ(*A*) − 1 decreases and approaches a lower asymptote when extreme epistasis effectively creates a single partially dominant locus (Figure 5). Selection for migration reduction increases with *ϕ*, because the dispersal modifier can reduce a larger fraction of the load when there are fewer loci contributing to local adaptation (as in Figure 2).

**Figure 5:**
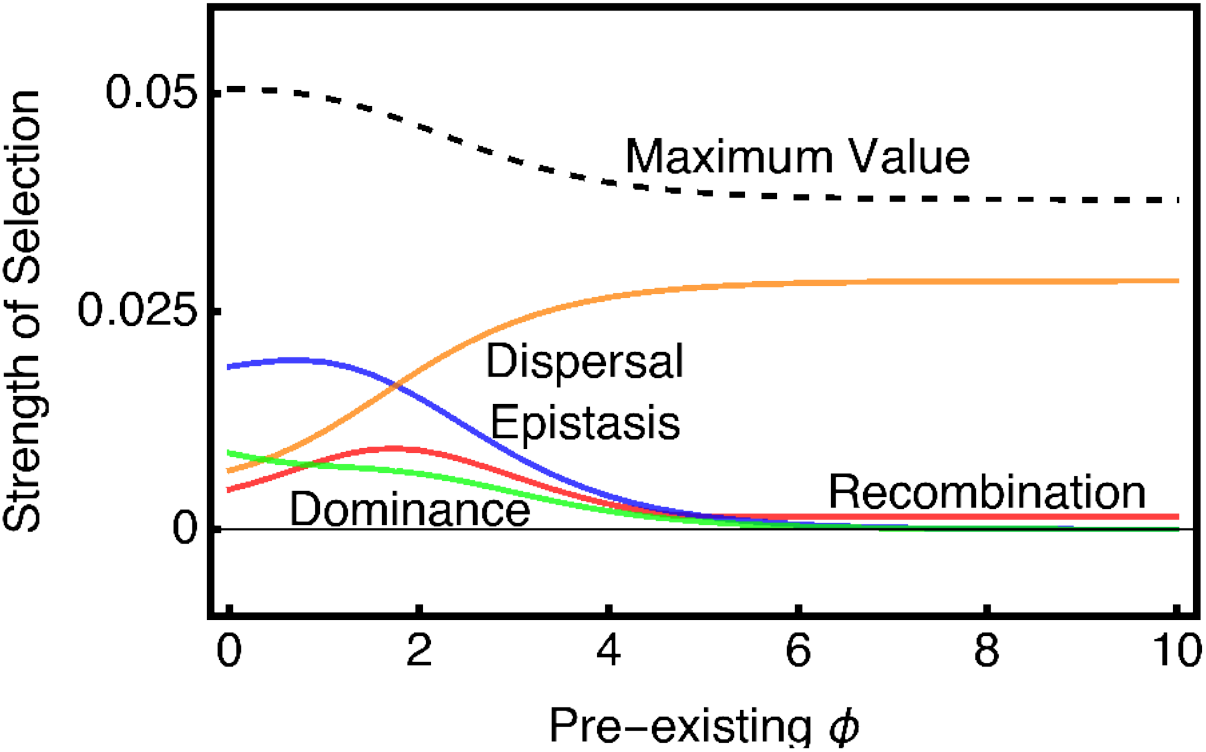
Strength of selection on different modifiers depends on the level of pre-existing epistasis. Parameter values for the ancestral population are *k* = 5, *S* = 0.1, *M* = 0.05, *R* = 0.5; *θ* = 0. *ϕ* is varied between 0 and 10. Strength of selection is shown for modifiers with extreme effects.

Selection for additional increases in epistasis shows a complex pattern. First, as *ϕ* increases from 0, there is an increase in selection for further increases in *ϕ*. Selection for further increases in *ϕ* then reaches a local maximum, and after *ϕ* reaches about 3 then selection for further increases in *ϕ* substantially decreases. Eventually, further selection for increased *ϕ* is reduced to zero. When *ϕ* = 0, selection for a complete reduction in recombination is about 1/4 as large as selection for increased *ϕ*.

Like selection for epistasis, recombination is initially more strongly selected as *ϕ* increases, and does not substantially decrease until *ϕ* is around 4. Selection decreases to about 4% of the theoretical maximum, but does not decay towards 0. Once *ϕ* is about about 4.5, selection for tight linkage supersedes selection for increased epistasis.

In contrast to the other modifier types, selection for dominance decreases as *ϕ* increases from 0. Once moderate levels of *ϕ* are reached, selection for tight linkage supersedes selection for dominance.

### Ancestral local adaptation

Ancestral levels of local adaptation will also determine heterozygosity and linkage disequilibrium necessary to generate selection on the modifiers. We thus also explored varying the parameter *S* which determines the fitness cost that the most maladapted genotype experiences. This analysis further allows us consider selection on the modifier under “weak” vs “strong” selection for local adaptation.

Figure 6A shows that selection on each type of modifier increases as the fitness benefit of local adaptation goes up. While the response of selection for dispersal modifiers appears linear, the other types of modifier show non-linear functions under stronger and stronger selection. However, the relative advantage of epistasis and dominance modifiers, as compared with recombination modifiers, remains large and constant from weak to strong selection (Figure 6B).

**Figure 6:**
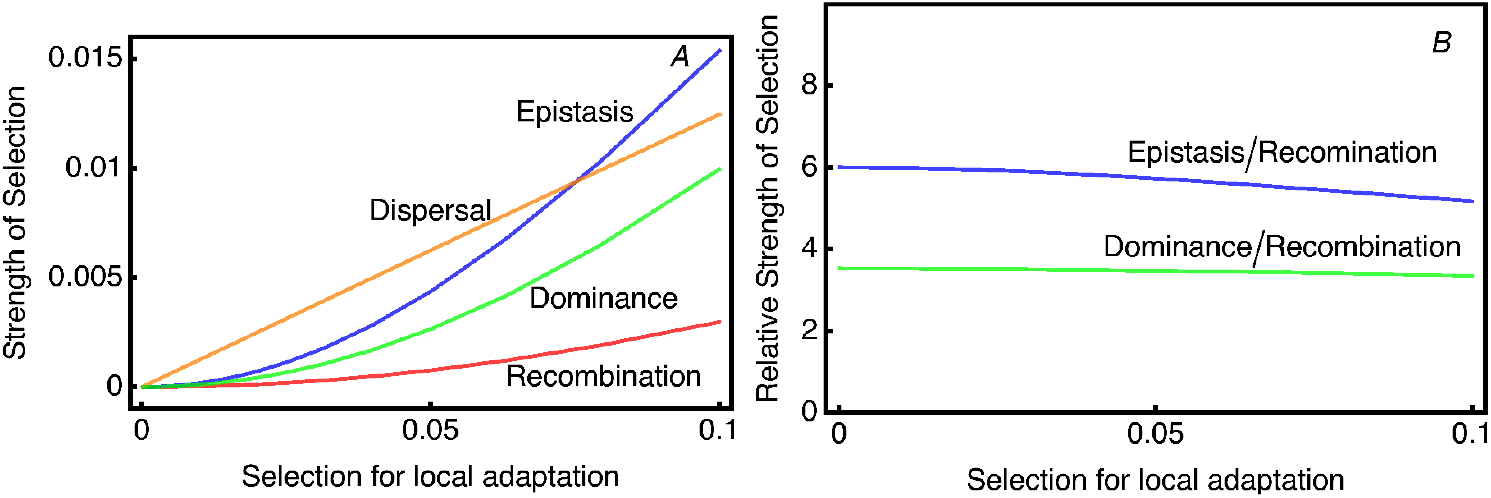
Selection for modifiers depends on the strength of selection for local adaptation. As the value of *S* is increased, the strength of selection on all types of modifier goes up (panel A). The relative advantage of epistasis and dominance modifiers, as compared with recombination modifiers, is shown in panel B. Epistasis modifiers have about a 6 fold higher invasion rate than a modifier that eliminates recombination (i.e. an inversion). Parameter values for the ancestral population are *k* = 3, *M* = 0.05, *R* = 0.5; *ϕ* = 0; *θ* = 0.

## Conclusions

We have presented a framework to compare the magnitude of selection between modifiers of the genetic architecture of a structured population undergoing local adaptation in two habitats with different environmental conditions. We identify four previously unreported features of selection on genetic architecture modifiers. First, selection for reduced recombination, even when the modifier is a chromosomal inversion that reduces recombination to zero, is weaker than for the other types of modifier. Second, when there is a constant total benefit of local adaptation but varying number of loci, increasing the number of loci increases selection strength for modifiers of epistasis and recombination, but decreases selection strength for dominance and migration modifiers. Third, for modifiers of dominance, the strength of selection is higher when they alter dominance at loci that the modifier itself is not linked to. Finally, even at moderate levels of epistasis in the resident population, selection for further increases in epistasis is stronger than selection for tighter linkage of the locally adapted loci. Selection on modifiers to reduce dispersal rates is complex, depending on the number of loci, epistasis and local adaptation of the ancestral population. Further study of this process would require building models that include selection for dispersal due to resource competition, source sink dynamics, or include a behavioral model for how short and long range dispersal are related.

Under migration selection balance when local populations are well mixed, then selection should act to reduce dispersal (Altenberg et al., 2017). Our results for selection to decrease dispersal are in line with this in that we see consistent directional selection towards lower dispersal rates. However, we find that selection on dispersal is weaker than on other types of modifiers unless the number of local adaptation loci is low and ancestral epistasis is strong. Adding complication, it is known that other factors, such as kin selection, might play a more important role in causing selection on dispersal than levels of migration load. When local resource competition between kin is possible, relatively high levels of dispersal are often favored (Hamilton and M., 1977; Ronce, 2007; Billiard and Lenormand, 2005). There is ample selection for dispersal to find a suitable habitat when local population density approaches the carrying capacity, and dispersal to find suitable habitat in these circumstances may lead to by-product dispersal at different distance scales. While migration itself is responsible for the entire fitness cost that drives selection for all other modifiers here investigated, selection acts more strongly on modifiers of genetic architecture than on modifiers of the dispersal rate. This seems paradoxical, because if selection acted to reduce dispersal then selection for other genetic architecture modifiers would disappear. How can we reconcile these ideas? Part of the answer is that modifiers of dispersal only alter the rate at which individuals carrying the modifier move between habitats, and do not change the impact or rate of interaction with other genotypes that have dispersed into a local deme. Modifiers that decrease dispersal protect the individual from arriving in a habitat to which they are not adapted, but it does not protect them from the effects of mating with others who themselves dispersed. Selection for reduced migration weakens as the number of loci increase because the immediate cost to a modifier of mating with maladapted individuals decreases.

Charlesworth and Barton (2018) point out that the strength of selection for inversions under continent-island selection is mathematically related to the amount of migration load generated in the absence of an inversion and the migration load generated once an inversion is fixed. Selection on an inversion, however, is determined by the fraction of the migration load that the inversion is able to ameliorate (Proulx and Phillips, 2005). Charlesworth and Barton (2018) consider a scenario where there is one-way migration from a continent to an island, fitness is multiplicative (i.e. additive on log scale), and locally adapted alleles are segregating on the island at migration-selection balance. An inversion that causes locally adapted alleles to become tightly linked can spread because it causes the marginal fitness of the haplotype carrying the modifier and a full complement of locally adapted alleles to be pegged at 1, and therefore is positively selected with a magnitude of *L* − *m*, or the load minus the migration rate (Appendix B). Under bi-directional reciprocal dispersal we consistently find that selection to reduce the recombination rate is weaker than other changes to the genetic architecture. This can further be analyzed by considering how the modifier alters the reproductive value classes of the offspring (Proulx and Servedio, 2009). We observe that when the modifier only partially reduces recombination then selection is quite weak.

Perhaps the most significant result of our analysis is that the strength of selection for dominance and epistasis modifiers is relatively stronger than that for recombination modifiers. This is in contrast to situations where locally adapted alleles are not yet established, linkage between the modifier and existing adaptation loci is important (Pontz and Bürger, 2022). We further found that modifiers of dominance spread fastest when they are unlinked to the locus they modify, but are linked to another locus that also contributes to local adaptation. To better appreciate this, consider two loci, A and B, both varying for alleles that confer local adaptation between habitats. If a modifier of dominance that causes the habitat 1 allele to be dominant at locus B/b and is in tight linkage with B, then it will confer a direct advantage whenever the modifier and B are in habitat 1 but a cost when in habitat 2. If the modifier is linked to the A allele at the A/a locus, then in habitat 1 it is beneficial when the carrier is heterozygous at the B/b locus. If the modifier migrates into the habitat two, it is only costly if one of the B alleles is not locally adapted. We thus expect that genomic islands of differentiation between populations are more likely caused by selection on partially-dominant mutations that just happen to appear next to already existing locally-adapted alleles. A possible example of this process might have occurred at the self-incompatibility locus of Brassica, where a non-recombining gene cluster showing high levels of DNA sequence divergence within and between species shows a hierarchy of partial dominance in locally-adapted populations (Durand et al., 2014).

In the work of Otto and Bourget (1999) an approximation is derived for the spread of a dominance modifier when there is one locus contributing to local adaptation. Otto and Bourget (1999) find that selection on a modifier can be approximated by the weighted average of the affect on heterozygote fitness in the two demes, where the weighting is the product of heterozygote frequencies and the reproductive value of alleles in each deme. This result reflects the selective value of changing a life-history parameter is the average over changes in the product of the immediate fitness effect of the mutation with the frequency of the class experiencing the change and the reproductive value of that class. In our study, heterozygote frequency alone is not proportional to the strength of selection on the modifier (results not shown), and this could be because changes in the ecological parameters, such as the dispersal rate, also cause the reproductive value of heterozygotes to change. Further work is required to relate our selection strength estimates to the underlying components of the frequency and reproductive value of the haplotypes.

Analogous to selection on recombination modifiers, selection for assortative mating preference can occur when heterozygotes are maintained by migration-selection balance, frequency dependence, or overdominance, and the effective recombination rate between locally-adapted loci can evolve, e.g., (Otto et al., 2008; Proulx and Servedio, 2009; Durinx and Van Dooren, 2009). Mating preference modifiers affect associations of genes without altering direct fitness, particularly for strong assortative mating, and their spread can be wholly driven by selection created by a migration load. Durinx and Van Dooren (2009) in particular have found that when load is caused by heterozygote advantage then modifiers of dominance are selected for, and also that assortative mating is selected for. When the frequency of heterozygotes is high, then selection for assortative mating can be stronger than selection for dominance. Our results here show that dominance modifiers can achieve selection coefficients of about one-third the maximum possible selection on modifiers, which means other types of modifier, such as mating preference modifiers, could also achieve substantially higher selection.

Most analyses of selection on modifiers assume that the fitness differences between alternative genotypes are small. There are two main reasons for this, one is analytical practicality and the other is the belief that the bulk of naturally segregating alleles have small effects on fitness. For the cases we studied here, both of these reasons are less applicable. Under bi-directional migration we do not have good analytical approximations for either the amount of migration load or the equilibrium genotype frequencies, so no reduction of mathematical complexity is achieved by assuming weak selection. We can still approximate selection on modifiers using a weak-load and quasi linkage equilibrium assumptions, and while these approximations perform well when numerically solving for the genotype frequency, they do not provide much additional insight (see Appendix B). Second, large differences in fitness between demes can be maintained with only a few locally-adapted loci and high dispersal rates, patterns which have been observed in natural populations, e.g. (Hoekstra et al., 2005; Hall and Willis, 2006). Our results indicate that modifiers of epistasis or dominance consistently experience larger selection coefficients than modifiers of recombination for both weak and strong migration loads.

In sum, we find little evidence for selection for decreased recombination rates under migration load. Instead, selection should diminish migration loads by favoring modifiers of dominance and epistasis within and between locally-adapted loci. Selection on dispersal modifiers is more complex. Our modeling assumes deterministic dynamics so a venue for future study is to find out if genetic drift and demographic stochasticity qualitatively change our conclusions.

## Acknowledgements

We thank Reinhard Bürger, Joachim Hermisson, Thomas Lenormand and Denis Roze for discussion, special thanks to Nick Barton for suggesting the form for the fitness function, and two anonymous reviewers for very helpful suggestions on how to improve the manuscript. HT is supported by the Agence Nationale pour la Recherche (ANR-17-CE02-0017-01, ANR-18-CE02-0017-01).

## Appendix A: Alternative life-cycle algebra

The life-cycle of the ancestral resident population can be described using tensor notation. This framework is useful as it can be modified to include complexities, such as non-Mendelian inheritance, transgenerational carryover environmental effects, etc., which the recursion equations presented in the main text cannot.

Given that there are *k* bi-allelic loci and random mating, the 2^*k*^ possible haplotypes are followed. The allele values are defined as 0/1 and the haplotypes are indexed by their binary numerical value offset by one, such that 0, 0, 0 maps to 1, and 0,0,1 maps to 2, etc. Haploid population frequency in deme *j* is presented as:

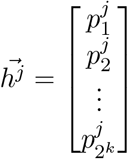

where 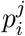 represents the frequency of the ith haplotype in the *j*th deme.

The diploid population frequency matrix is then formed by taking the outer product of 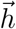 with itself, i.e. multiplying the frequency of all haplotype pairs in all possible orders:

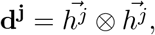

with migration then mixing the densities between demes:

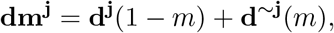

where ∼ *j* inverts the binary value of *j*, and therefore represents the index for the other deme.

Diploid individual fitness matrix is defined as 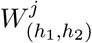 is the fitness of an (*h*_1_, *h*_2_) individual in deme *j* and has dimension [2^*k*^, 2^*k*^]:

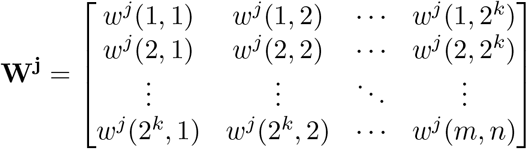

After selection, we re-normalize the vector of genotype frequencies by dividing by population mean fitness which ensures that the genotype frequencies sum to 1.

The diploid frequency after migration and selection is then:

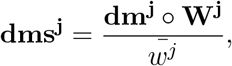

where we use to represent element-by-element matrix multiplication (i.e. the Hadamard product); the mean fitness in deme *j* being 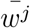.

The next stage of the life-cycle is gamete production where meiotic segregation and recombination of parental haplotypes takes place. Gamete production is accomplished by Hadamard tensor multiplication, which means that each diploid genotype frequency is multiplied by the vector of recombination probabilities of producing each gamete haplotype. For example, in a one locus scenario the gamete production vector is v = [1/2, 1/2], so if the frequency of a heterozygote is *x* then the product is *x* ∗ *v* = [*x*/2, *x*/2]. The gamete production tensor is a rank 3 tensor of length 2^*k*^ in each dimension, giving

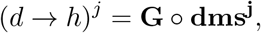

where now 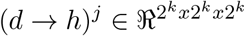 where 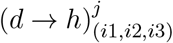 represents the frequency of type *i* 3 haplotypes produced by parents with diploid genotype (*i*1, *i*2). While (*i*1, *i*2) and (*i*2, *i*1) parents are identical in all ways, they appear as separate entries for computational convenience. And the total frequency of haplotypes is found by summing up the terms for a specific haplotype, that is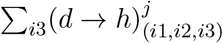. This can be represented in tensor operations by first unfolding, or flattening, (*d* → *h*)^*j*^ by a single level and then by taking the inner product with a one’s vector.

Having described the life-cycle of the resident population we compare the notation presented in the main text with the tensor notation above. The ancestral population will attain a population genetic equilibrium between migration, selection, mating, segregation, recombination and syngamy, after iterating:

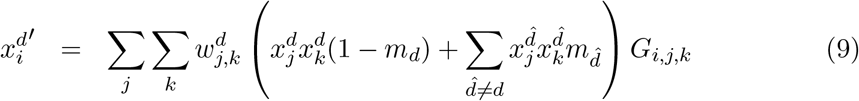

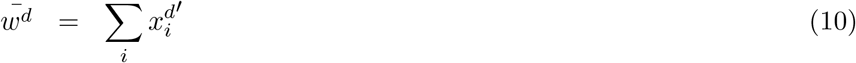

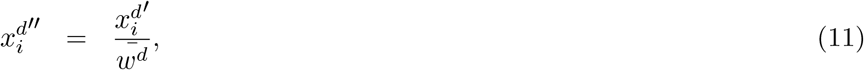

where 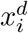 is the frequency of haplotype *i* in deme *d*, 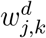 is the fitness of genotype (*j, k*) in deme *d, G*_*i,j,k*_ represents the probability that an adult with haplotypes *j* and *k* will produce a gamete of haplotype *i*, and 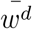 is the average fitness within deme *d*. Haplotype frequency in the next generation is given by 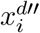.

With tensor notation these recursion equations can be represented as:

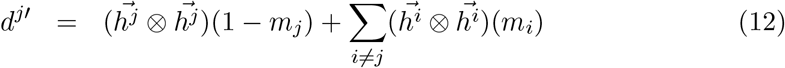

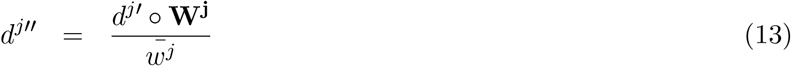

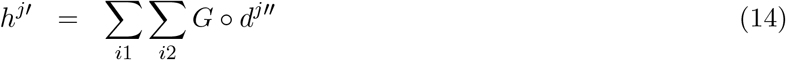

where equations 9, 10, 11, are analogous to equations 12, 13, 14 respectively.

## Appendix B: Inversions and epistasis modifiers

We wish to understand the spread of inversions, eliminating recombination between local adaptation loci, as a function of selection on modifiers of epistasis. For this, we first write several definitions to simplify notation.

The frequency of haplotypes carrying *i* locally maladapted alleles is in deme *d* defined as *x*_(*d*)(*i*)_. Define the average fitness among haplotypes that do not disperse (residents) from deme *d* as 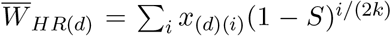, and the average fitness among haplotypes that do disperse (migrants) as 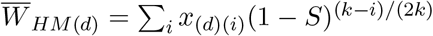. The diploid average fitness is in deme *d* is 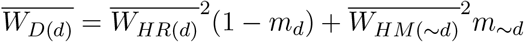where ∼ *d* represents “not deme d”, i.e. the other deme. We then define the fitness load among non-dispersing haplotypes as 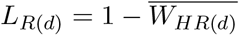, and the fitness load among dispersing haplotypes as 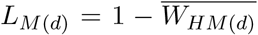. The total haploid load is defined as 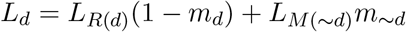.

The invasion matrix for the inversion is given by

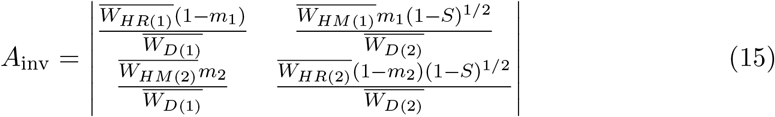

Define λ_inv_ as the dominant eigenvalue of *A*_inv_ which is the discrete time growth factor, i.e. *R*_0_, for the inversion.

With unidirectional migration from deme 2 to 1, the matrix reduces to a lower diagonal matrix with an eigenvalue of 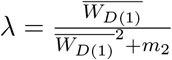. We define the selection coefficient for the inversion as s_inv_ = λ_inv_ − 1, which is approximately *s*_inv_ = *L* − *m*, which is equation 4 from Charlesworth and Barton (2018). Note that this equation is quite general, it does not depend on any assumptions about the number of loci. It stems from the simple fact that the inversion haplotype has constant marginal fitness set to 1 and that the frequency of the inversion haplotype is reduced by *m* each generations (see also Proulx and Phillips (2005) for a general derivation of this result).

Under bi-directional migration with symmetry between the demes, we can find the eigenvalues and then approximate around weak selection. Under symmetry, we suppress the deme indices. We first Taylor expand around weak load, and then Taylor expand around small *S*, including second order terms (supplementary Mathematica file). We found that our approximation was both qualitatively and quantitatively accurate when we include the second order terms in S, but fails qualitatively when we only include the first order terms. Defining *s*_inv_ = λ − 1, we have

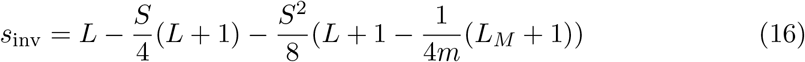

It is difficult to gain much intuition directly from this expression.

We develop a similar approximation for a complete modifier of epistasis, using the simplifying assumption that the modifier is always in a haplotype that contains at least one allele that is locally adapted in deme 1. This gives the matrix:

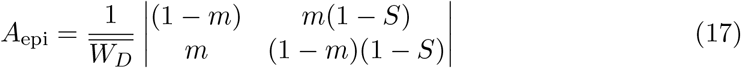

Approximating around weak load and weak *S*, and dropping terms of order *S*^2^ ∗ *L* we come to:

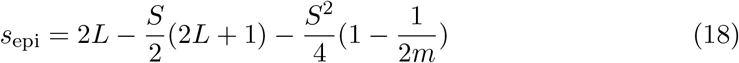

## Legends

**S1 Mathematica Wolfram Language Script file**. **Methods Code**. This file contains code to create the recombination tensors, to perform the resident population numerical simulations, and to create the functions that create the modifier invasion matrix. This file is stored in the Wolfram Language Script file, which can be read into a regular Mathematica session. This file must be loaded (<< Methods.wls) in order to run the code to create the figures.

**S2 Mathematica Wolfram Language MX file**. **Executable Invasion Matrix Functions**. This file contains binary executable code to create the invasion matrices for the modifier allele. These functions can be loaded (<<InvaderFunctions.mx) in order to create the matrices.

**S3 Mathematica Notebook. Compiled Figures**. This is a Mathematica Note-book file that contains code to load the methods and create the figures. All of the figures in the paper can be created with this code.

